# Transmembrane channel-like proteins regulate crop size and contraction dynamics during *Drosophila* feeding

**DOI:** 10.1101/2025.04.07.647529

**Authors:** Dharmendra Kumar Nath, Subash Dhakal, Youngseok Lee

## Abstract

Feeding regulation in *Drosophila* involves complex interactions between mechanosensory and neuroendocrine pathways. Our study identified transmembrane channel-like (TMC) proteins as key regulators of crop size and contraction, functioning through distinct neuronal populations. They influence crop size via diuretic hormone, *Dh44*-expressing neuroendocrine cells in the pars intercerebralis (PI) region and regulate crop contraction through the serotonin receptor 5-HT7. We found that TMC proteins are broadly expressed from the gut to the brain, reinforcing their role in the brain-gut axis. Mechanotransduction channels, including NompC, Piezo, and TMC, facilitate food ingestion, with TMC channels playing an additional role in food storage and transport. We noted the coexpression of *piezo* with *Dh44* in only two neurons, indicating that at least two *Dh44* cells are required for crop size regulation. Moreover, we identified Dh44R2 as the key receptor regulating crop size. Unlike DH44 and Piezo, TMC, 5-HT7, and TRPγ are essential for crop contraction, suggesting that these channels serve as therapeutic targets for regulating food intake. Our findings also support the involvement of a mechanosensory serotonergic pathway in regulating crop physiology, integrating sensory and neuroendocrine signals to control food storage and transport. These findings advance our understanding of the neuronal and molecular mechanisms underlying feeding behavior in *Drosophila* and provide a foundation for exploring conserved pathways that regulate food intake in other organisms, including mammals.

## INTRODUCTION

The fruit fly *Drosophila melanogaster* is a powerful model organism for studying genetics, development, and physiology. Among its many biological processes, feeding behavior is essential for survival; this encompasses food detection, ingestion, storage, and digestion, in addition to nutrient absorption (Volkoff and Rønnestad, 2020). The gastrointestinal (GI) tract of *Drosophila* is structurally divided into three main regions: the foregut, midgut, and hindgut (Gartner, 1985; King, 1988). The anterior foregut consists of the esophagus, crop, and cardia. The crop is a unique diverticulated organ present only in the order Diptera. It plays a key role in food storage, early digestion, detoxification, and microbial control (Edgecomb et al., 1994; Miguel-Aliaga et al., 2018). A complex network of valves and sphincters regulates the movement of food into and out of the crop before it enters the main alimentary canal. After ingestion, food passes through the pharynx and is transported down the esophagus via peristalsis before being stored in the crop. Rhythmic contractions of the proventriculus then propel the food into the midgut for digestion and nutrient absorption. The midgut is connected to the hindgut, where waste products from the Malpighian tubules mix with undigested material before excretion. Malpighian tubules, which are blind-ended ducts, play a key role in excretion and osmoregulation (Chopra et al., 2022).

In insects, nutrient regulation is controlled by food intake, gut motility, digestive enzyme secretion, and absorption, which are modulated by neuropeptides in the stomatogastric nervous system. These neuropeptides can stimulate or inhibit enzymatic activity in the gut, thereby affecting digestion (Fadda et al., 2019; Harshini et al., 2002). The central nervous system (CNS) and sensory nervous system (SNS) innervate the gut and mouthparts of insects (Spit et al., 2012) while enteric neurons play a crucial role in intestinal transit and peristalsis regulation. Muscle valves within each gut region are innervated by neurons that generate peristaltic waves, directly linking peristalsis to nutrient demand (Cognigni et al., 2011). Enteric neurons also regulate food ingestion volume (Olds and Xu, 2014).

Several neuropeptides, including serotonin, adipokinetic hormone (Akh), myosuppressin, allatostatin (AST), diuretic hormone 44 (DH44), FMRFamide, and short neuropeptide F (sNPF), have been identified in the *Drosophila* midgut (Chopra et al., 2022; Sano et al., 2015). Notably, *Drosophila* Piezo neurons, which innervate the anterior gut and crop, regulate food ingestion volume (Min et al., 2021). Piezo ion channels, which regulate mechanotransduction, are involved in various physiological processes, including nephrocyte function, cardiac performance, stomach stretch, and defecation (Koehler and Denholm, 2021; Landino, 2023; Oh et al., 2021; Puri et al., 2025; Zechini et al., 2022). Similarly, transmembrane channel-like (TMC) proteins act as mechanosensors in proprioception and hearing (Guo et al., 2016; Kawashima et al., 2015). These proteins are evolutionarily conserved and contribute to sensory functions across species; they play a role in body position detection, movement coordination, auditory processing, and substrate hardness discrimination in *Drosophila* (Guo et al., 2016; He et al., 2019; Yue et al., 2019).

Although crop expansion has been well characterized, the mechanisms underlying crop contraction remain poorly understood. The assessment of crop contraction in *D. melanogaster* will provide a unique opportunity to explore the regulation of nutrient storage and utilization. The findings may also have broader implications for understanding metabolism and growth regulation in multicellular organisms. In this study, we identified a functional role of *tmc* in regulating crop size and contraction. We found that *tmc^1^*mutants exhibit an enlarged crop and increased crop contraction rate. Our results indicate that *tmc*-mediated crop size regulation occurs via Dh44 neuroendocrine cells in the brain, while crop contraction is regulated through the serotonin receptor 5-hydroxytryptamine 7 (5-HT7).

## RESULTS

### *tmc^1^* mutants exhibit enlarged crop size and increased crop contractions

To determine whether the physiological function of the crop involves mechanosensation, we analyzed crop size and contraction in wild-type (*w^1118^*) and *tmc^1^*male flies. We fed the flies a diet consisting of 1% agarose supplemented with 5% sucrose and blue dye for 12 h. We then captured the images under normal conditions and assessed crop size after dissection. We observed that *tmc^1^*mutants exhibited a significantly larger crop size than control flies (Fig. 1A–D). To quantify crop size (Dhakal et al., 2022) we established a scoring system ranging from 1 to 5 based on the amount of blue dye retained in the crop; a score of 1 indicated no crop expansion, while 5 indicated the largest crop expansion (Fig. 1E). Given the established role of Piezo (Qin et al., 2024) and TMC proteins in food swallowing, we measured both crop size and contraction in *w^1118^*, *piezo^KO^*, and *tmc^1^* mutant flies. Both *piezo^KO^* and *tmc^1^* mutant flies exhibited a significantly larger crop size than control flies (Fig. 1F). More than 60% of control flies exhibited an intermediate crop size with a score of 3 (70.55% ± 2.18%), with only a small proportion reaching a score of 4 (12.50% ± 2.57%) or 5 (5.83% ± 2.09%). In contrast, a significantly higher proportion of *piezo^KO^*and *tmc^1^* mutants exhibited crop size scores of 4 and 5. Specifically, crop size scores of 3, 4, and 5 were noted in 19.28% ± 3.29%, 36.51% ± 3.16%, and 27.30% ±1.71% of *tmc^1^* mutant flies, respectively, and in 26.67% ± 2.11%, 30.67% ± 1.63%, and 21.33% ± 2.49% of *piezo^KO^* flies, respectively.

**Figure 1.**
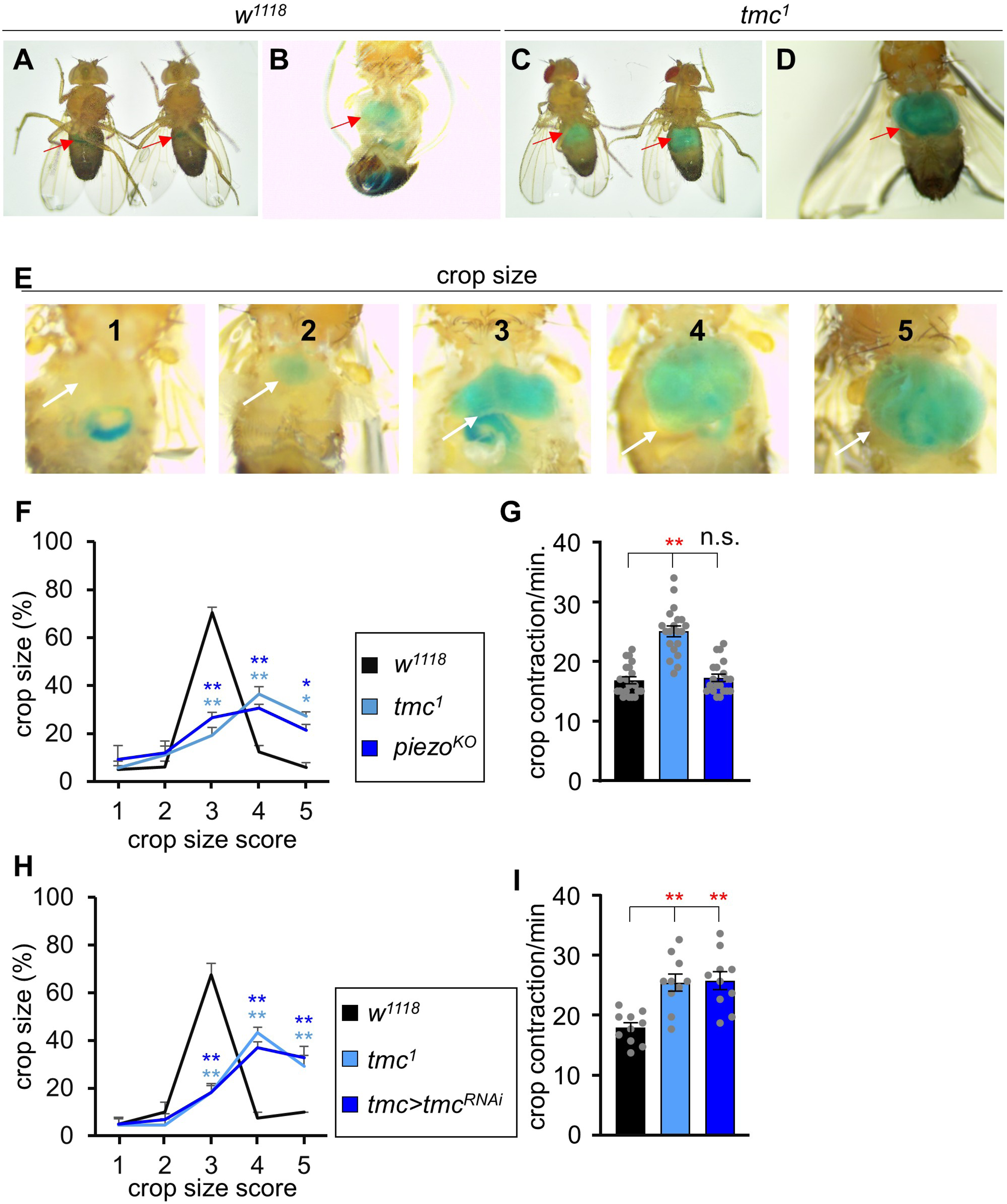
Larger crop size and higher crop contraction in *tmc^1^* mutants. **(A)** Representative images showing the crop size of *w^1118^* flies after feeding 1% agarose containing 5% sucrose and blue dye for 12 h under normal conditions. **(B)** Representative images showing the crop size of *w^1118^* flies after feeding 1% agarose containing 5% sucrose and blue dye for 12 h after dissection of the abdomen. **(C)** Representative images of the crop of *tmc^1^* flies after feeding 1% agarose containing 5% sucrose and blue dye for 12 h under normal conditions. **(D)** Representative images of the crop of *tmc^1^* flies after feeding 1% agarose containing 5% sucrose and blue dye for 12 h after dissection of the abdomen. **(E)** Crop size scores from 1 to 5 based on the crop size in *w^1118^* flies after feeding 1% agarose containing 5% sucrose and blue dye for 12 h. **(F)** Percentage of *w^1118^*, *piezo^KO^*, and *tmc^1^* flies with crop size scores ranging from 1 to 5 after feeding 1% agarose containing 5% sucrose and blue dye for 12 h. n = 5–6. **(G)** Crop contraction rate after feeding 1% agarose containing 5% sucrose and blue dye for 12 h. n = 20. **(H)** Percentage of *w^1118^*, *tmc^1^*, and *UAS*-*tmc^RNAi^*/*tmc*-*GAL4*;*UAS*-*dicer2*/+ flies with crop size scores ranging from 1 to 5 after feeding 1% agarose containing 5% sucrose and blue dye for 12 h. n = 4. **(I)** Crop contraction rate in *w^1118^*, *tmc^1^*, and *UAS*-*tmc^RNAi^*/*tmc*-*GAL4*;*UAS*-*dicer2*/+ flies after feeding 1% agarose containing 5% sucrose and blue dye for 12 h. n = 10. All values are reported as the means ± SEMs. Comparisons between multiple experimental groups were performed via single-factor ANOVA with Scheffe’s *post hoc* test. Asterisks indicate significant differences from the controls (\**P* < 0.05, \*\**P* < 0.01). Each dot represents the distribution of individual sample values.

Next, we measured crop contraction frequency (contractions/min). In control (*w^1118^*) flies, the average contraction rate was 16.85 ± 0.59 contractions/min (Fig. 1G). In contrast, *tmc^1^* mutants exhibited a significantly higher contraction rate of 25.05 ± 0.90 contractions/min. The contraction rate in *piezo^KO^* flies was comparable to that in controls (17.25 ± 0.62 contractions/min) (Fig. 1G). As a previous study implicated the involvement of nompC in food swelling (Qin et al., 2024) we also assessed crop size and contraction in nompC⁴ mutants. Both parameters appeared normal (Fig. EV1A,B), suggesting that nompC does not play a major role in crop function.

To rule out potential sex-specific effects, we assessed crop size and contraction in female flies. The results mirrored those observed in male flies (Fig. EV1C,D). Moreover, RNAi-mediated knockdown of *tmc* under the control of *tmc*-*GAL4* resulted in a phenotype similar to that of *tmc^1^*mutants, further supporting the role of TMC proteins in regulating crop size and contraction (Fig. 1H,I). These findings indicate that TMC proteins are required for proper crop function, as their loss results in defects in the regulation of crop size and contraction frequency.

### TMC proteins are essential for the regulation of crop size and contraction dynamics

To further confirm the role of TMC proteins in crop physiology, we inactivated *tmc* cells using the inwardly rectifying potassium channel *UAS*-*Kir2*.*1* (Hodge, 2009) and measured both crop size and contraction rate. We noted that *tmc*-*GAL4*-inactivated flies exhibited a significantly larger crop size and higher crop contraction rate than control *UAS*-*Kir2*.*1* flies (Fig. 2A,B). We also performed immunostaining to examine the expression pattern of TMC proteins in the crop. We detected TMC protein expression in key crop-associated structures, including the proventriculus, crop duct, and crop itself (Fig. 2C,D), suggesting a functional role of these proteins in crop physiology.

**Figure 2.**
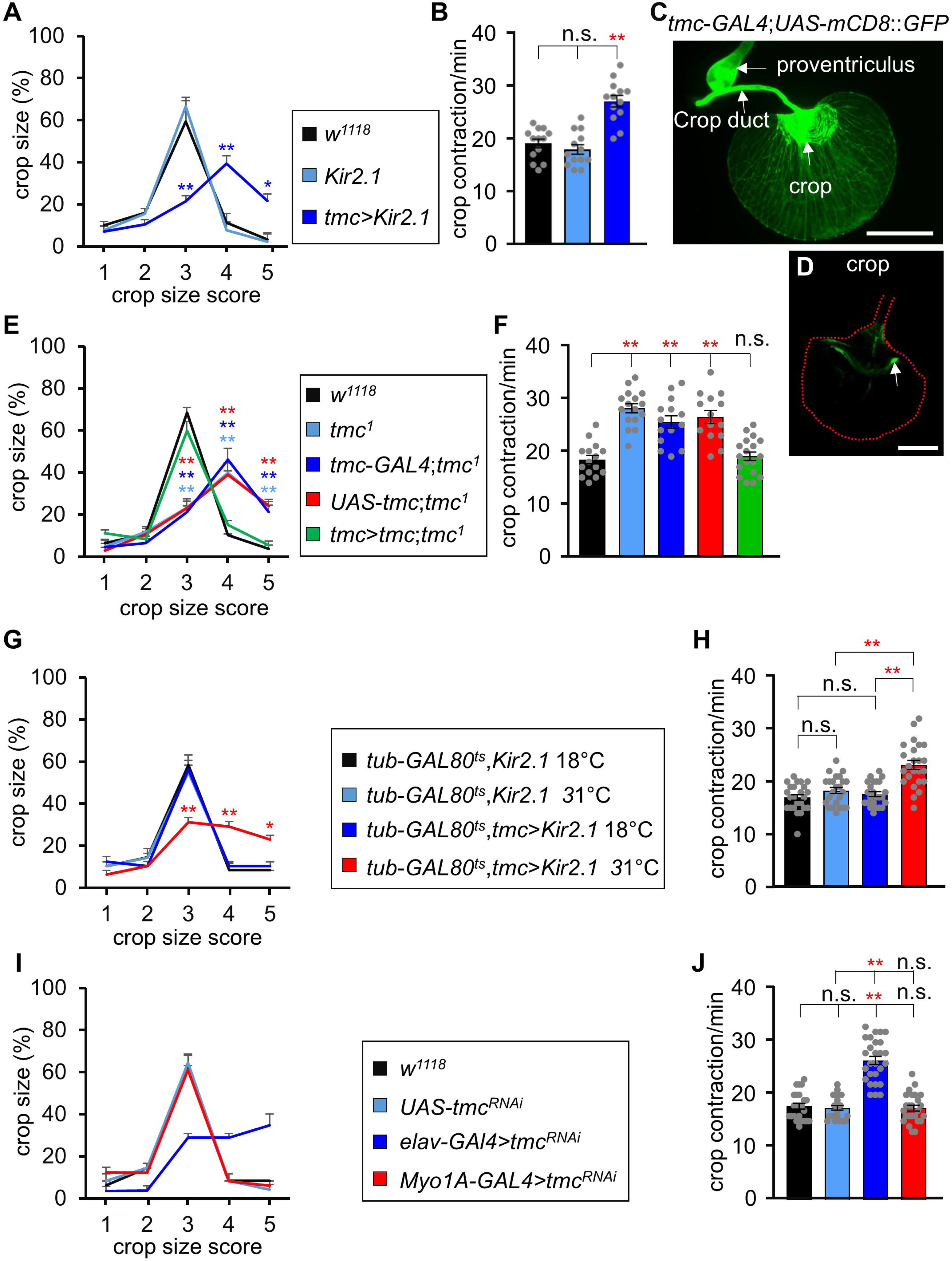
*tmc* is essential for the regulation of crop size and crop contraction. **(A)** Percentage of *w^1118^*, *UAS*-*Kir2*.*1*, and *tmc*-*GAL4*/*UAS*-*Kir2*.*1* flies with crop size scores ranging from 1 to 5 after feeding 1% agarose containing 5% sucrose and blue dye for 12 h. n = 5. **(B)** Crop contraction rate in *w^1118^*, *UAS*-*Kir2*.*1*, and *tmc*-*GAL4*/*UAS*-*Kir2*.*1* flies after feeding 1% agarose containing 5% sucrose and blue dye for 12 h. n = 13. (**C–D**) Images of *tmc* reporter (*tmc*-*GAL4*;*UAS*-*mCD8*::*GFP*) in the crop region. (**C**) Expression of *tmc* in the foregut region, including the proventriculus, crop duct, and crop (indicated with an arrow). Scale bar represents 50 µm. (**D**) Expression of *tmc* neurons in the crop. Scale bar represents 50 µm. **(E)** Percentage of *w^1118^*, *tmc^1^*, *tmc*-*GAL4*;*tmc^1^*, *UAS*-*tmc*;*tmc^1^*, and *tmc*-*GAL4*/*UAS*-*tmc*;*tmc^1^* flies with crop size scores ranging from 1 to 5 after feeding 1% agarose containing 5% sucrose and blue dye for 12 h. n = 4. **(F)** Crop contraction rate in *w^1118^*, *tmc^1^*, *tmc*-*GAL4*;*tmc^1^*, *UAS*-*tmc*;*tmc^1^*, and *tmc*-*GAL4*/*UAS*-*tmc*;*tmc*^1^ flies after feeding 1% agarose containing 5% sucrose and blue dye for 12 h. n = 14–18. **(G)** Percentage of *tub*-*GAL80^ts^*;*UAS*-*Kir2*.*1* and *tub*-*GAL80^ts^*/*tmc*-*GAL4*;*UAS*-*Kir2*.*1*/+ flies with crop size scores ranging from 1 to 5 after feeding 1% agarose containing 5% sucrose and blue dye for 12 h at 18°C and 31°C (permissive and nonpermissive temperatures for the temperature sensitive GAL4 repressor *tub*-*GAL80^ts^*, respectively). n = 4. **(H)** Crop contraction rate in *tub*-*GAL80^ts^*;*UAS*-*Kir2*.*1* and *tub*-*GAL80^ts^*/*tmc*-*GAL4*;*UAS*-*Kir2*.*1*/+ flies after feeding 1% agarose containing 5% sucrose and blue dye for 12 h at 18°C and 31°C (permissive and nonpermissive temperatures for the temperature-sensitive GAL4 repressor *tub*-*GAL80^ts^*, respectively). n = 24. **(I)** Percentage of *w^1118^*, *UAS*-*tmc^RNAi^*;*UAS*-*dicer2*, *UAS*-*tmc^RNAi^*/+;*elav*-*GAL4*/*UAS*-*dicer2*, and *Myo1A*-*GAL4*/*UAS*-*tmc^RNAi^*;*UAS*-*dicer2*/+ flies with crop size scores ranging from 1 to 5 after feeding 1% agarose containing 5% sucrose and blue dye for 12 h. n = 2–3. **(J)** Crop contraction rate in *w^1118^*, *UAS*-*tmc^RNAi^*;*UAS*-*dicer2*, *UAS*-*tmc^RNAi^*/+;*elav*-*GAL4*/*UAS*-*dicer2*, and *Myo1A*-*GAL4*/*UAS*-*tmc^RNAi^*;*UAS*-*dicer2*/+ flies after feeding 1% agarose containing 5% sucrose and blue dye for 12 h. n = 24. All values are reported as the means ± SEMs. Comparisons between multiple experimental groups were performed via single-factor ANOVA with Scheffe’s *post hoc* test. Asterisks indicate significant differences from the controls (\**P* < 0.05, \*\**P* < 0.01). Each dot represents the distribution of individual sample values.

To determine whether *tmc* is directly involved in crop regulation, we expressed *UAS*-*tmc* under the control of *tmc*-*GAL4*. This rescued the *tmc^1^* mutant phenotype, restoring the crop size and contraction rate to normal levels (Fig. 2E,F). Furthermore, to assess the developmental role of *tmc*-expressing cells in crop physiology, we performed a temperature-shift experiment using *tub*-*GAL80^ts^*;*UAS*-*Kir2*.*1*. Compared to controls, flies reared at 18°C and subsequently shifted to 31°C exhibited an altered crop size and increased contraction rate (Fig. 2G,H), indicating a dynamic regulatory role of *tmc*-expressing cells in crop function.

To determine whether TMC proteins function in neurons or gut cells, we knocked down *tmc* using *tmc^RNAi^* under the control of the pan-neuronal driver *elav*-*GAL4* and the gut enterocyte-specific driver *Myo1A*-*GAL4*. Knockdown via *elav*-*GAL4* phenocopied the crop size and contraction defects observed in *tmc^1^* mutants, whereas knockdown via *Myo1A*-*GAL4* resulted in no significant changes (Fig. 2I,J). Together, these findings highlight the crucial role of TMC proteins in regulating crop size and contraction dynamics, further supporting their involvement in regulating crop physiology.

### *Dh44* neurons regulate crop size through TMC-dependent mechanisms

Previously, our group demonstrated that *Dh44* signaling in the pars intercerebralis (PI) region of the brain regulates crop size (Dhakal et al., 2022). To further investigate the role of *Dh44* in crop size regulation, we inactivated *Dh44*-*GAL4* using *UAS*-*Kir2.1* and measured the crop size and contraction rate. The inactivation of *Dh44* neurons resulted in a higher proportion of flies with crop size scores of 4 and 5 and a lower proportion of flies with a crop size score of 3 (Fig. 3A). However, the crop contraction rate remained unchanged (Fig. 3B).

**Figure 3.**
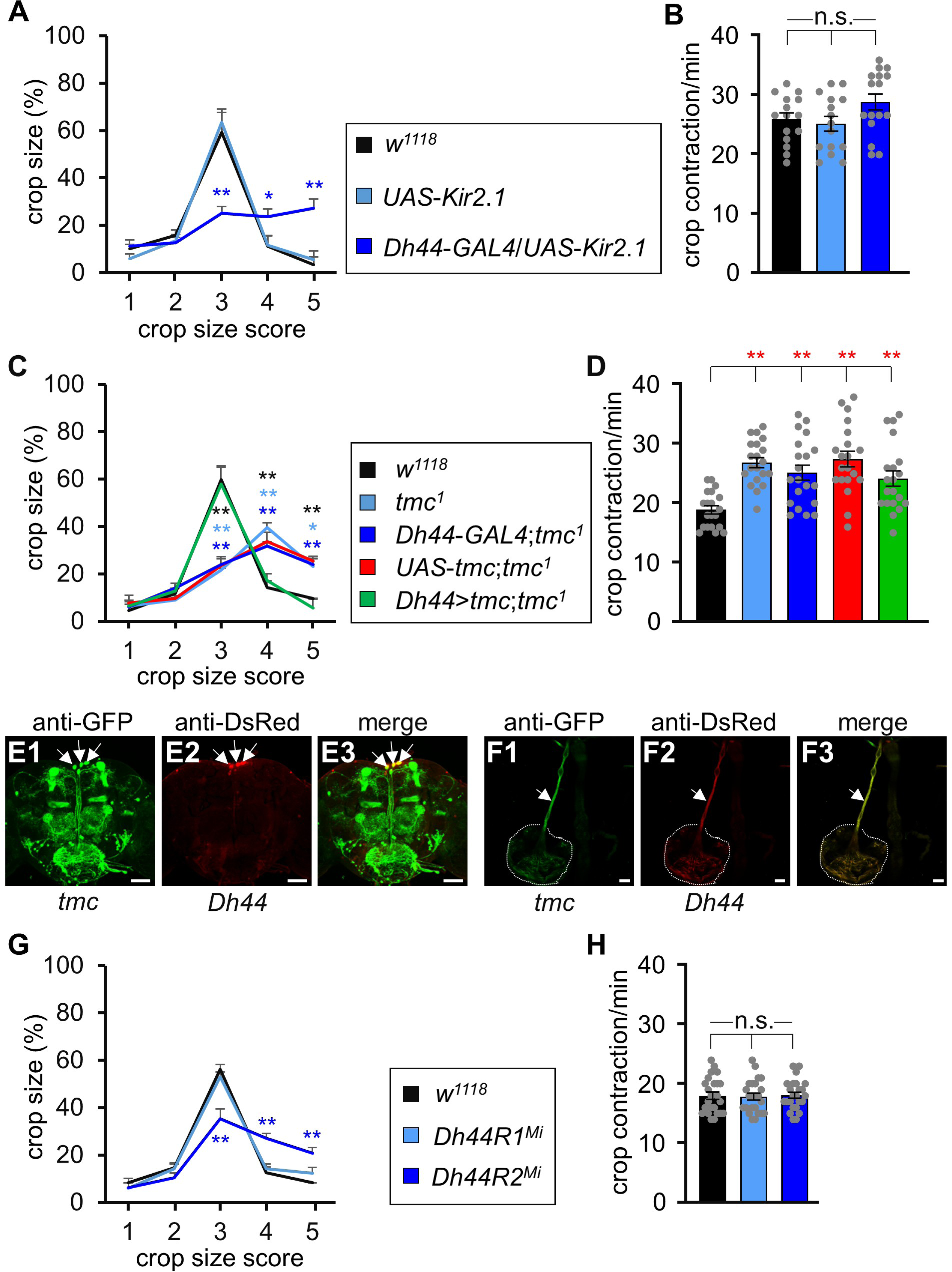
Involvement of *Dh44* in *tmc*-mediated crop size regulation. **(A)** Percentage of *w^1118^*, *UAS*-*Kir2*.*1*, and *Dh44*-*GAL4*/*UAS*-*Kir2*.*1* flies with crop size scores ranging from 1 to 5 after feeding 1% agarose containing 5% sucrose and blue dye for 12 h. n = 5. **(B)** Crop contraction rate in *w^1118^*, *UAS*-*Kir2*.*1*, and *Dh44*-*GAL4*/*UAS*-*Kir2*.1 flies after feeding 1% agarose containing 5% sucrose and blue dye for 12 h. n = 15–16. **(C)** Percentage of *w^1118^*, *tmc^1^*, *Dh44*-*GAL4*;*tmc^1^*, *UAS*-*tmc*;*tmc^1^*, and *Dh44*-*GAL4*/*UAS*-*tmc*;*tmc^1^* flies with crop size scores ranging from 1 to 5 after feeding 1% agarose containing 5% sucrose and blue dye for 12 h. n = 5. **(D)** Crop contraction rate in *w^1118^*, *tmc^1^*, *tmc*-*GAL4*;*tmc^1^*, *UAS*-*tmc*;*tmc^1^*, and *tmc*-*GAL4*/*UAS*-*tmc*;*tmc^1^* flies after feeding 1% agarose containing 5% sucrose and blue dye for 12 h. n = 20. (**E1–E3**) Confocal images showing the coexpression of *tmc* and *Dh44* (*Dh44*-*LexA*/*tmc*-*GAL4*;*LexAop*-*mCherry*/*UAS-mCD8*::*GFP*) in the PI region of an adult fly based on immunohistochemical analysis. Scale bar represents 50 µm. (**F1–F3**) Confocal images showing the expression of *tmc* and *Dh44* (*Dh44*-*LexA*/*tmc*-*GAL4*;*LexAop*-*mCherry*/*UAS-mCD8*::*GFP*) in the crop and crop duct of an adult fly based on immunohistochemical analysis. Scale bar represents 50 µm. **(G)** Percentage of *w^1118^*, *Dh44R1^Mi^*, and *Dh44R2^Mi^* flies with crop size scores ranging from 1 to 5 after feeding 1% agarose containing 5% sucrose and blue dye for 12 h. n = 4. **(H)** Crop contraction rate in *w^1118^*, *Dh44R1^Mi^*, and *Dh44R2^Mi^*flies after feeding 1% agarose containing 5% sucrose and blue dye for 12 h. n = 24. All values are reported as the means ± SEMs. Comparisons between multiple experimental groups were performed via single-factor ANOVA with Scheffe’s *post hoc* test. Asterisks indicate significant differences from the controls (\**P* < 0.05, \*\**P* < 0.01). Each dot represents the distribution of individual sample values.

Next, we expressed *UAS*-*tmc* under the control of *Dh44*-*GAL4* and assessed its effect on crop physiology. *UAS*-*tmc* expression under the control of *Dh44*-*GAL4* successfully rescued the crop size defect observed in *tmc^1^* mutants (Fig. 3C); however, the crop contraction rate remained unaffected (Fig. 3D). We then performed immunostaining of the brain to determine whether *tmc* and *Dh44* neurons are coexpressed. We found that three *tmc*-expressing neurons colocalized with *Dh44* neurons in the PI region (Fig. 3E1–E3). We also examined *tmc* and *Dh44* expression in the crop duct (Fig. 3F1–F3). Our group previously reported that *Dh44^Mi^* mutant exhibit an enlarged crop size but normal crop contraction rate (Dhakal et al., 2022). To further understand the role of DH44 signaling, we measured crop size and contraction in flies lacking DH44 receptors (*Dh44R1^Mi^* and *Dh44R2^Mi^*). Although *Dh44R1^Mi^* mutants exhibited a normal crop size, *Dh44R2^Mi^* mutants exhibited increased crop size scores of 4 and 5 (Fig. 3G). However, the crop contraction rate remained normal in both *Dh44R1^Mi^* and *Dh44R2^Mi^* mutants (Fig. 3H).

These findings suggest that TMC-mediated crop size regulation occurs via *Dh44*-expressing neurons in the brain, highlighting a neural mechanism linking *Dh44* signaling to crop function.

### Piezo plays a role in *Dh44* neurons

In *Drosophila*, Piezo plays a crucial role in regulating food consumption volume (Min et al., 2021). To assess the interaction between TMC and Piezo proteins in crop physiology, we expressed *UAS*-*tmc* under the control of *piezo*-*GAL4* and measured the crop size and contraction rate (Fig. 4A,B). *UAS*-*tmc* expression under the control of *piezo*-*GAL4* successfully restored the increased crop sizes (scores 4 and 5) in *tmc^1^*mutants to normal levels (Fig. 4A). However, the crop contraction rate remained significantly higher than that in control flies (Fig. 4B).

**Figure 4.**
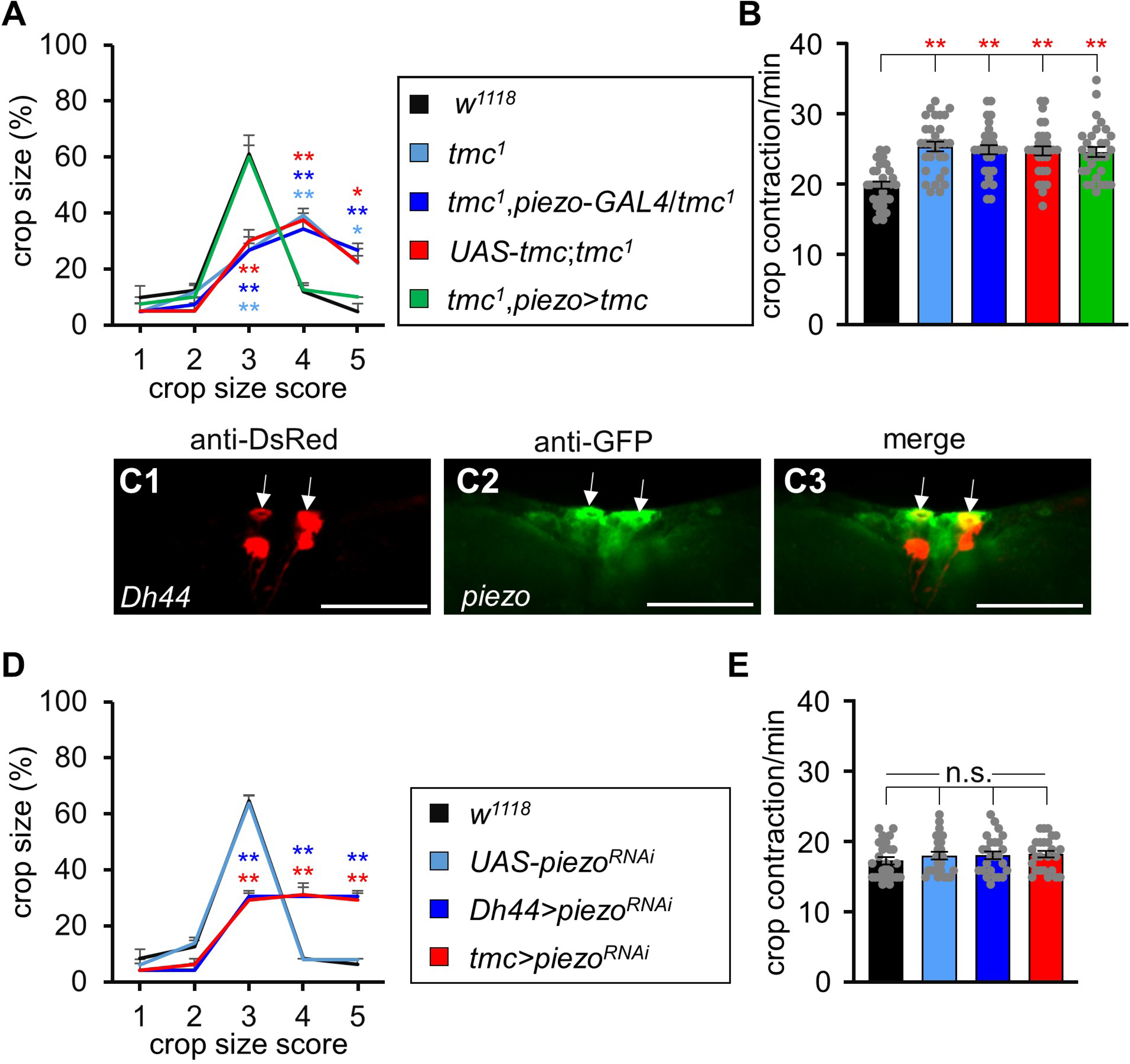
Recapitulation of crop size defects in *tmc^1^* mutants using *piezo*-*GAL4*. **(A)** Percentage of *w^1118^*, *tmc^1^*, *tmc^1^*,*piezo*-*GAL4*/+, *UAS*-*tmc*;*tmc^1^*, and *UAS*-*tmc*/+;*tmc^1^*,*piezo*-*GAL4* flies with crop size scores ranging from 1 to 5 after feeding 1% agarose containing 5% sucrose and blue dye for 12 h. n = 4. **(B)** Crop contraction rate in *w^1118^*, *tmc^1^*, *tmc^1^*,*piezo*-*GAL4*/+, *UAS*-*tmc*;*tmc^1^*, and *UAS*-*tmc*/+;*tmc^1^*,*piezo*-*GAL4* flies after feeding 1% agarose containing 5% sucrose and blue dye for 12 h. n = 30. (**C1–C3**) Confocal images showing the coexpression of *Dh44* and *piezo* (*Dh44*-*LexA*/*UAS*-*mCD8*::*GFP*;*LexAop*-*mCherry*/*piezo*-*GAL4*) in the PI region of the brain. Scale bar represents 100 µm. **(D)** Percentage of *w^1118^*, *UAS*-*piezo^RNAi^*, *UAS*-*piezo^RNAi^*/+;*UAS*-*dicer2*/+,*Dh44*-*GAL4*/+, and *piezo^RNAi^*/+,*tmc*-*GAL4*/+,*UAS*-*dicer2*/+ flies with crop size scores ranging from 1 to 5 after feeding 1% agarose containing 5% sucrose and blue dye for 12 h. n = 3–4. **(E)** Crop contraction rate in *w^1118^*, *UAS*-*piezo^RNAi^*, *UAS*-*piezo^RNAi^*/+;*UAS*-*dicer2*/+,*Dh44*-*GAL4*/+, and *piezo^RNAi^*/+,*tmc*-*GAL4*/+,*UAS*-*dicer2*/+ flies after feeding 1% agarose containing 5% sucrose and blue dye for 12 h. n = 24–25. All values are reported as the means ± SEMs. Comparisons between multiple experimental groups were performed via single-factor ANOVA with Scheffe’s *post hoc* test. Asterisks indicate significant differences from the controls (\**P* < 0.05, \*\**P* < 0.01). Each dot represents the distribution of individual sample values.

As *Dh44* neurons play a role in crop regulation, we examined potential interactions between *piezo* and *Dh44* neurons in the brain. Immunostaining revealed that two *piezo*-expressing neurons were coexpressed with *Dh44* neurons in the PI region of the brain (Fig. 4C1–C3). Moreover, we knocked down *piezo^RNAi^* in *Dh44*-*GAL4* and *tmc*-*GAL4*. The knockdown of *piezo^RNAi^* under the control of *Dh44*-*GAL4* and *tmc*-*GAL4* increased the crop size scores to 4 and 5; however, the crop contraction frequency remained normal (Fig. 4D,E). These findings further support the role of TMC proteins in regulating crop size through *Dh44* neurons, reinforcing a mechanosensory pathway involving Piezo, TMC, and DH44 signaling in the regulation of crop physiology.

### Serotonin modulates TMC-dependent regulation of crop size and contraction

Serotonin (5-HT) regulates gut motility and is primarily produced by enterochromaffin (EC) cells in the GI tract of mammals (Najjar et al., 2023). While serotonin is widely recognized for its role in the CNS, the majority of it is produced in the gut, where it regulates various physiological processes, including gut motility (Martin et al., 2022; Najjar et al., 2023). During peristalsis, serotonin is released to enhance gut motility (Tanveer et al., 2024). Peripheral serotonin also plays a key role in regulating gut motility (Zhang et al., 2024).

In insects, the innervation of serotonergic neurons in the crop and gut regions suggests their involvement in food transport along the digestive tract (Moffett and Moffett, 2005; Molaei and Lange, 2003). To further understand the regulatory role of TMC proteins in crop physiology, we performed *tmc^RNAi^* knockdown in five different serotonin receptor gene GAL4 lines, including *5-HT1A*, *5-HT1B*, *5-HT2A*, *5-HT2B*, and *5-HT7*. The knockdown of *tmc^RNAi^* under the control of *5-HT7*-*GAL4* recapitulated the *tmc^1^* crop phenotype, resulting in an enlarged crop size and increased crop contraction rate, while the other receptors appeared to be dispensable (Fig. 5A,B).

**Figure 5.**
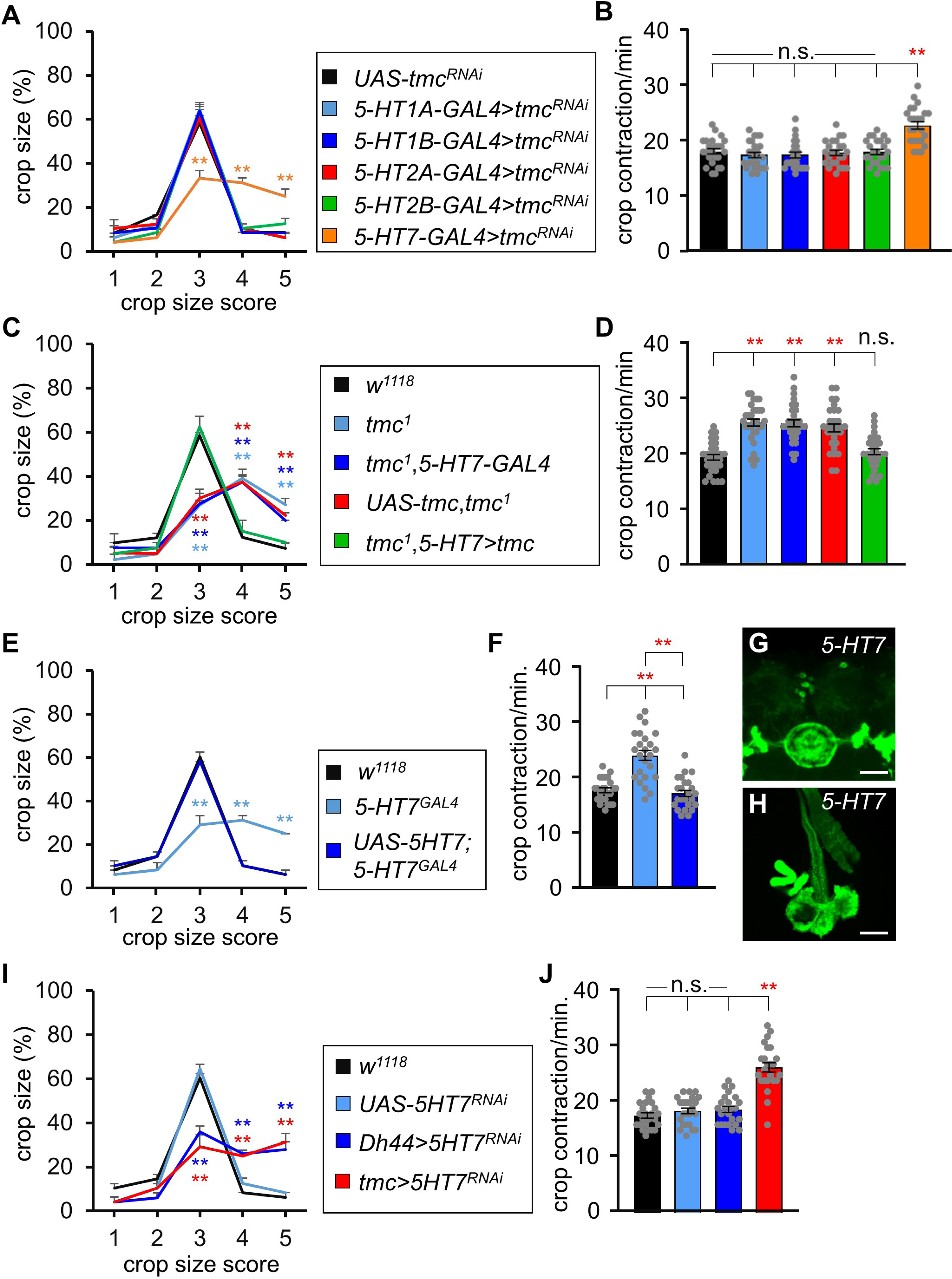
Role of 5-HT7 in TMC-mediated regulation of crop size and contraction. **(A)** Percentage of *5*-*HT1A^GAL4^*, *5*-*HT1B*-*GAL4*, *5*-*HT2A*-*GAL4*, *5*-*HT2B*-*GAL4*, and *5*-*HT7^GAL4^*flies with crop size scores ranging from 1 to 5 after feeding 1% agarose containing 5% sucrose and blue dye for 12 h after the knockdown of *UAS*-*tmc^RNAi^*. n = 4. **(B)** Crop contraction rate in *5*-*HT1A^GAL4^*, *5*-*HT1B*-*GAL4*, *5*-*HT2A*-*GAL4*, *5*-*HT2B*-*GAL4*, and *5*-*HT7^GAL4^* flies after feeding 1% agarose containing 5% sucrose and blue dye for 12 h after the knockdown of *UAS*-*tmc^RNAi^*. n = 24. **(C)** Percentage of *w^1118^*, *tmc^1^*, *tmc^1^*,*5*-*HT7^GAL4^*/*tmc^1^*, *UAS*-*tmc*;*tmc^1^*, and *UAS*-*tmc*/+;*tmc^1^*/*tmc^1^*,*5*-*HT7^GAL4^* flies with crop size scores ranging from 1 to 5 after feeding 1% agarose containing 5% sucrose and blue dye for 12 h. n = 4. **(D)** Crop contraction rate in *w^1118^*, *tmc^1^*,*5*-*HT7^GAL4^*/*tmc^1^*, *UAS*-*tmc*;*tmc^1^*, and *UAS*-*tmc*/+;*tmc^1^*/*tmc^1^*,*5*-*HT7^GAL4^*flies after feeding 1% agarose containing 5% sucrose and blue dye for 12 h. n = 30. **(E)** Percentage of *w^1118^*, *5*-*HT7^GAL4^*, and *UAS*-*5HT7*;*5*-*HT7^GAL4^* flies with crop size scores ranging from 1 to 5 after feeding 1% agarose containing 5% sucrose and blue dye for 12 h. n = 4. **(F)** Crop contraction rate in *w^1118^*, *5*-*HT7^GAL4^*, and *UAS*-*5HT7*;*5*-*HT7^GAL4^*flies after feeding 1% agarose containing 5% sucrose and blue dye for 12 h. n = 24. **(G)** Confocal image showing the expression of *5*-*HT7^GAL4^*(*UAS*-*mCD8*::*GFP*;*5*-*HT7^GAL4^*) in the brain. Scale bar represents 100 µm. **(H)** Confocal image showing the expression of *5*-*HT7^GAL4^* (*UAS*-*mCD8*::*GFP*;*5*-*HT7^GAL4^*) in the crop. Scale bar represents 100 µm. **(I)** Percentage of *w^1118^*, *UAS*-*5HT7^RNAi^*, *UAS*-*dicer2*/+;*Dh44*-*GAL4*/*UAS*-*5HT7^RNAi^*, and *tmc*-*GAL4*/+;*UAS*-*dicer2*/*UAS*-*5HT7^RNAi^* flies with crop size scores ranging from 1 to 5 after feeding 1% agarose containing 5% sucrose and blue dye for 12 h. n = 4. **(J)** Crop contraction rate in *w^1118^*, *UAS*-*5HT7^RNAi^*, *UAS*-*dicer2*/+;*Dh44*-*GAL4*/*UAS*-*5HT7^RNAi^*, and *tmc*-*GAL4*/+;*UAS*-*dicer2*/*UAS*-*5HT7^RNAi^*flies after feeding 1% agarose containing 5% sucrose and blue dye for 12 h. n = 24. All values are reported as the means ± SEMs. Comparisons between multiple experimental groups were performed via single-factor ANOVA with Scheffe’s *post hoc* test. Asterisks indicate significant differences from the controls (\*\**P* < 0.01). Each dot represents the distribution of individual sample values.

To assess whether *5-HT7*-*GAL4* can rescue the *tmc^1^*crop phenotype, we expressed *UAS*-*tmc* under its control. This successfully restored the crop size defects (Fig. 5C) and crop contraction rate (Fig. 5D) observed in *tmc^1^* mutants to normal level. We also assessed crop regulation using *5-HT7* knock-in *GAL4* (*5-HT7^GAL4^*) mutants and found that these mutants exhibited similar crop defects to *tmc^1^*mutants (Fig. 5E,F). To further confirm the functional role of *5-HT7* in regulating crop size and contraction, we expressed *UAS-5-HT7* under the control of *5-HT7^GAL4^*; this successfully restored the crop size and contraction rate to normal levels (Fig. 5E,F). We then examined 5-HT7 expression in the PI region of the brain and the crop (Fig. 5G,H) and provided anatomical evidence for its role in crop physiology. To explore potential coordination among *5-HT7*, *Dh44*, and *tmc* for crop size regulation, we knocked down *5-HT7* using *5*-*HT7^RNAi^*under the control of *Dh44*-*GAL4* and *tmc-GAL4* and then measured the crop size and contraction rate (Fig. 5I,J). The knockdown of *5*-*HT7^RNAi^*under the control of *Dh44*-*GAL4* increased the crop size; however, the contraction rate remained unaffected. This finding suggests that 5-HT7 regulates crop size through DH44 signaling but does not influence contraction dynamics. In contrast, the knockdown of *5*-*HT7^RNAi^*under the control of *tmc*-*GAL4* increased the crop size and contraction rate.

These findings suggest that serotonin-specific 5-HT7 signaling plays a key role in TMC-mediated regulation of crop size and contraction, further reinforcing the interplay among serotonin, DH44, and TMC proteins for regulating crop physiology.

### Food texture affects crop contraction rate but does not correlate with crop size

Food attributes, such as texture, nutritional value, and taste, significantly influence food preference. Studies have revealed that the mechanosensory neuron *tmc* plays a functional role in food texture discrimination (Zhang et al., 2016). In this study, we investigated whether the hardness of food affects its transport from the crop to the midgut during digestion.

We fed *w^1118^*, *piezo^KO^*, and *tmc^1^*mutant flies food containing 1%–5% agar with 5% sucrose and blue dye and measured the crop size and contraction rate. We found that the crop size of the flies remained unchanged regardless of food hardness (Fig. 6A–C). In contrast, the crop contraction rate significantly decreased with an increase in agar concentration in *w^1118^* and *piezo^KO^* flies. However, *tmc^1^*mutants maintained a consistent contraction rate (Fig. 6D). These findings suggest that food hardness influences the crop contraction rate; however, the crop size remains unaffected.

**Figure 6.**
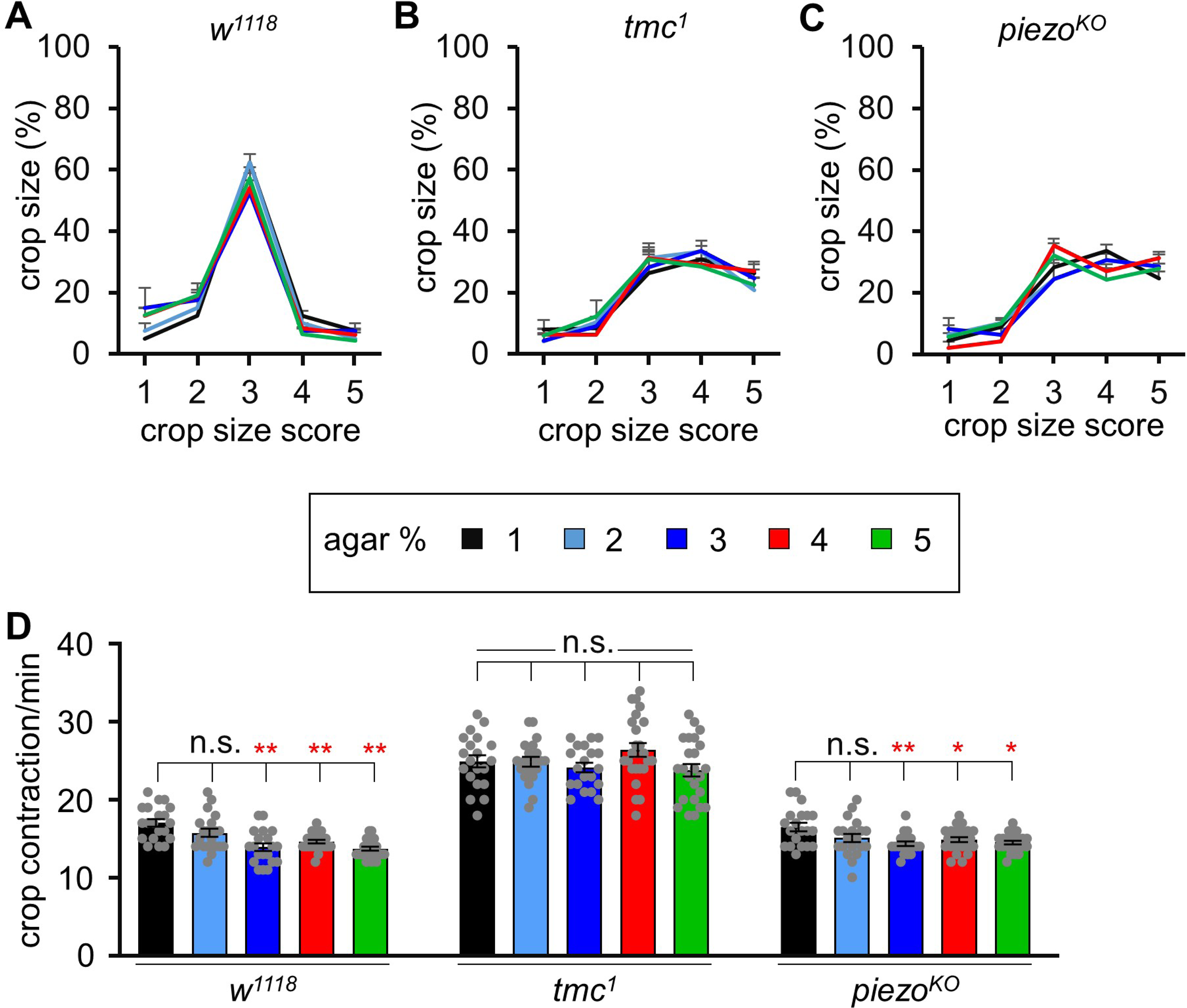
Hardness of food affects the crop contraction rate. **(A)** Percentage of *w^1118^* flies with different crop size scores after feeding different concentrations of agarose (1%–5%) containing 5% sucrose and blue dye for 12 h. n = 4. **(B)** Percentage of *tmc^1^* flies with different crop size scores after feeding different concentrations of agarose (1%–5%) containing 5% sucrose and blue dye for 12 h. n = 4. **(C)** Percentage of *piezo^KO^* flies with different crop size scores after feeding different concentrations of agarose (1%–5%) containing 5% sucrose and blue dye for 12 h. n = 4. **(D)** Crop contraction rate in *w^1118^*, *tmc^1^*, and *piezo^KO^* flies after feeding agarose (1%– 5%) containing 5% sucrose and blue dye for 12 h. n = 20. All values are reported as the means ± SEMs. Comparisons between multiple experimental groups were performed via single-factor ANOVA with Scheffe’s *post hoc* test. Asterisks indicate significant differences from the controls (\**P* < 0.05, \*\**P* < 0.01). Each dot represents the distribution of individual sample values.

## Discussion

### Neuronal regulation of crop size during feeding behavior

In *Drosophila*, the CNS regulates energy homeostasis and modulates feeding behavior based on internal states (Pool and Scott, 2014). The regulation of food intake involves the integration of multisensory information and various neuromodulatory systems. Several signaling pathways, including those involving orthologs of mammalian peptidergic signals, such as tachykinin, cholecystokinin, neuropeptides, and insulin, have been functionally conserved throughout evolution (Al-Anzi et al., 2010; Fleischer et al., 2018; Söderberg et al., 2012). In animals, neurons in the brain integrate sensory input with internal hunger and satiety cues to control feeding behavior (Cifuentes and Acosta, 2022). We found that TMC proteins are essential for regulating both crop size and contraction rate, with distinct cell populations mediating these physiological processes.

The fly proboscis contains *tmc*-expressing cells that can detect and respond to specific tastants based on texture, triggering signaling cascades that influence feeding behavior (Zhang et al., 2016). Three mechanotransduction channels, namely NompC, Piezo, and TMC, coordinately function in food swallowing and are linked to brain motor neurons (Qin et al., 2024). Our findings extend this role, demonstrating that TMC proteins function in a later stage of food processing, including food storage in the crop and transport from the crop to the midgut. The widespread expression of TMC-positive neurons throughout the brain, from the PI region to the subesophageal zone (SEZ), suggests its involvement in multiple neuronal circuits.

Crop size is directly linked to food ingestion volume; greater ingestion leads to larger crop expansion, a process primarily regulated by *Dh44*-expressing cells in the PI region. The expression pattern of *tmc* from the gut to the brain highlights its role in the brain-gut axis. Previously, we found that the transient receptor potential channel γ (TRPγ) regulates feeding behavior and crop physiology via *Dh44* neuroendocrine cells (Dhakal et al., 2022). Similarly, in this study, we found that *tmc*-mediated crop size regulation occurs via *Dh44* cells. DH44 is a key neuroendocrine hormone involved in metabolic regulation, physiological homeostasis, and ion balance (Cabrero et al., 2002). In *Drosophila*, *Dh44* plays diverse roles in feeding behavior, acting as a hunger signal, a nutrient sensor, a metabolic regulator, a feeding rhythm controller, and an integrator of internal and external cues (Dhakal et al., 2022; Nath et al., 2024; Oh et al., 2021). Of the six *Dh44*-expressing neurons in the PI region—a known neuroendocrine center, (Dus et al., 2015)—three are coexpressed with *tmc*, suggesting a significant role of *tmc* in regulating food ingestion. On the other hand, *piezo* is coexpressed with *Dh44* in only two neurons, indicating that at least two *Dh44* neurons are required to regulate crop size based on food intake levels. Further research on the precise mechanisms underlying *Dh44*-mediated feeding regulation will provide deeper insights into the neuronal circuits and signaling pathways governing food intake in flies and potentially other organisms.

We also unraveled a novel role of 5-HT7 in regulating crop size through its function in *Dh44* cells. Moreover, we identified that only one *Dh44* receptor, *Dh44R2*, is involved in crop size regulation, whereas *Dh44R1* appears dispensable for this function.

### Serotonergic control of crop contraction

Serotonin plays a well-established role in regulating feeding behavior across various invertebrates. It acts as a neurotransmitter and a neuromodulator, coordinating feeding activity (Orchard, 2006). In *Drosophila*, various neurotransmitters, such as dopamine and serotonin, modulate feeding drive and satiety (Neckameyer and Bhatt, 2012; Yapici et al., 2016). Our findings demonstrate that *tmc*-mediated crop contraction occurs via the serotonin receptor 5-HT7. This aligns with prior evidence suggesting that serotonin is a crucial regulator of feeding behavior that influences hunger, satiety, and mood-related feeding aspects through both central and peripheral mechanisms (Voigt and Fink, 2015) (Song and Avery, 2012). Moreover, serotonin, in combination with corticotropin-releasing factor (CRF)-related peptides, can regulate diuresis in some insects, such as *Rhodnius prolixus*, implicating its involvement in the termination of rapid post-feeding diuresis (Pool and Scott, 2014). A deeper understanding of the complex roles of serotonin in feeding regulation could provide insights into the neural circuits and molecular pathways underlying hunger control and may have implications for addressing eating disorders in humans.

Importantly, our study clarifies the distinct roles of TMC, 5-HT7, and TRPγ in controlling crop contraction rates. On the other hand, Dh44, Dh44R2, and Piezo play no significant role in this process. Given the influence of these mechanosensory and serotonergic pathways on feeding regulation, TMC, 5-HT7, and TRPγ may serve as potential targets for developing therapeutic strategies to control food intake in humans. Future studies in mammals will be required to validate this possibility.

Overall, our findings highlight the crucial role of TMC proteins in the feeding process. These proteins coordinate with the neuroendocrine hormone *Dh44* in the PI region of the brain to regulate crop size, while they interact with the serotonin receptor 5-HT7 to modulate crop contraction. Crop contraction is also influenced by the texture of ingested food, independent of overall crop size. These findings advance our understanding of the neuronal and mechanosensory mechanisms governing feeding behavior and serve as a foundation for future research on the conserved regulatory pathways controlling food intake.

## Materials and Methods

### Reagents and tools table

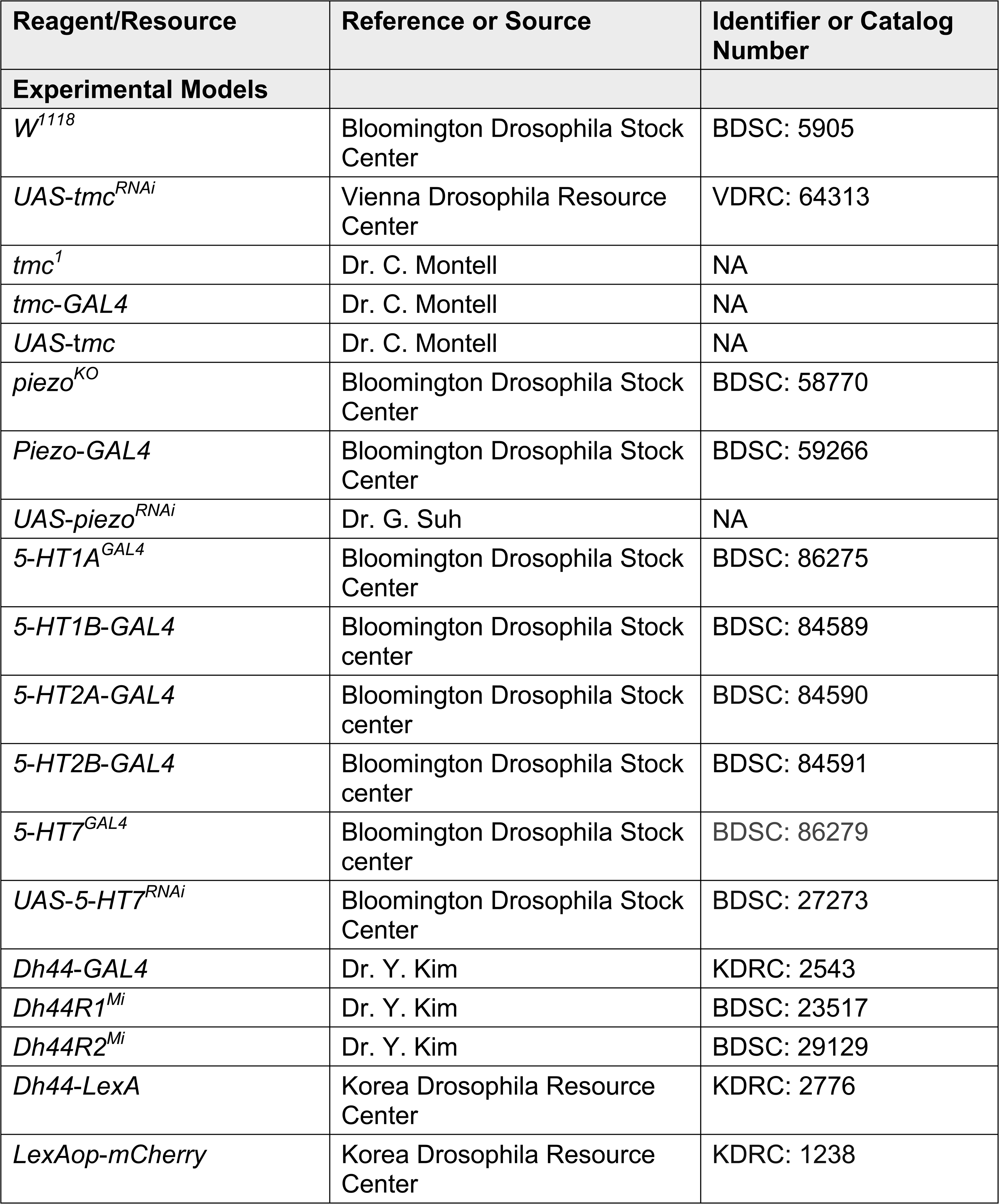
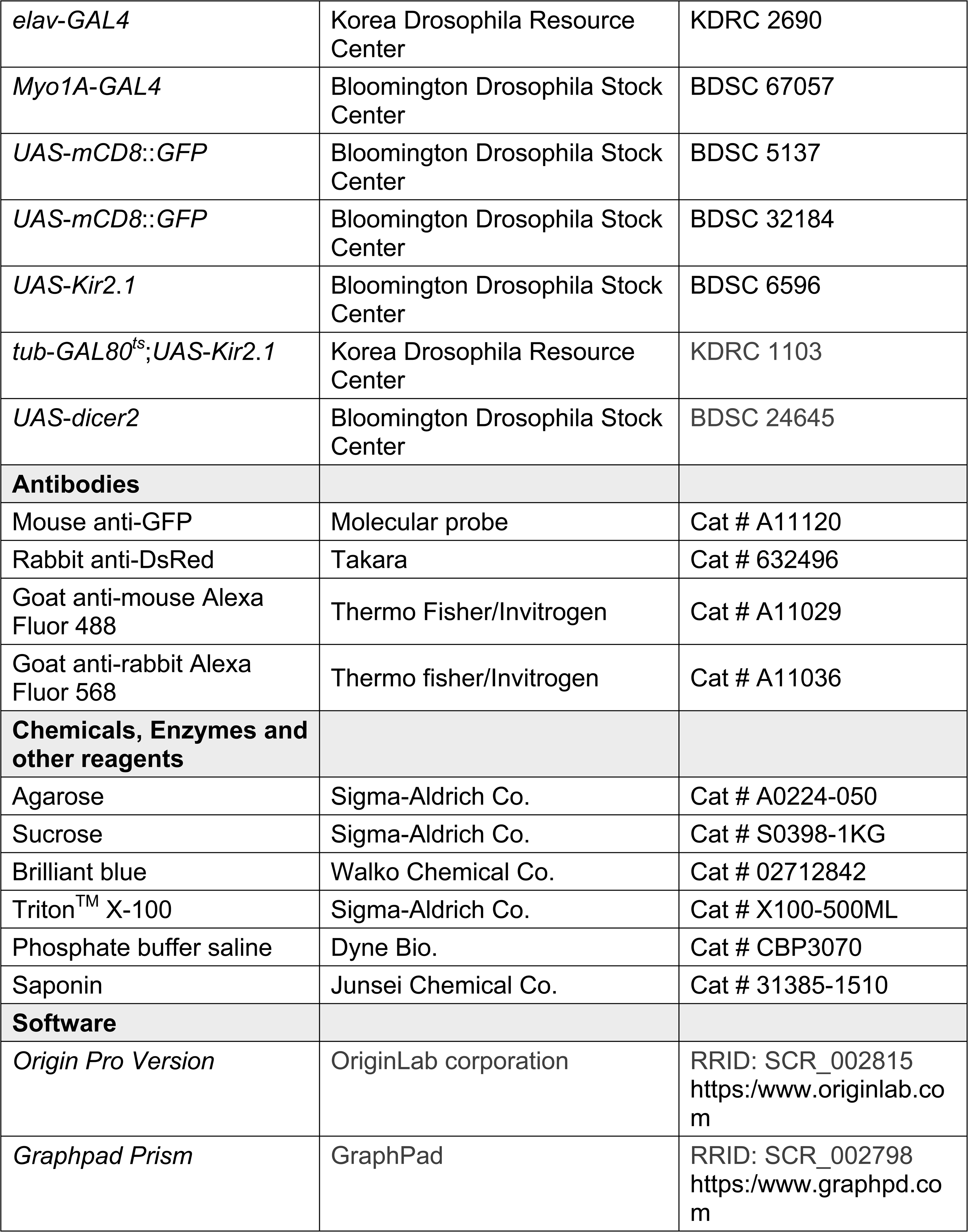

| Reagent/Resource | Reference or Source | Identifier or Catalog Number |
| --- | --- | --- |
| **Experimental Models** |  |  |
| *W<sup>1118</sup>* | Bloomington Drosophila Stock Center | BDSC: 5905 |
| *UAS-tmc<sup>RNAi</sup>* | Vienna Drosophila Resource Center | VDRC: 64313 |
| *tmc<sup>1</sup>* | Dr. C. Montell | NA |
| *tmc-GAL4* | Dr. C. Montell | NA |
| *UAS-tmc* | Dr. C. Montell | NA |
| *piezo<sup>KO</sup>* | Bloomington Drosophila Stock Center | BDSC: 58770 |
| *Piezo-GAL4* | Bloomington Drosophila Stock Center | BDSC: 59266 |
| *UAS-piezo<sup>RNAi</sup>* | Dr. G. Suh | NA |
| *5-HT1A<sup>GAL4</sup>* | Bloomington Drosophila Stock Center | BDSC: 86275 |
| *5-HT1B-GAL4* | Bloomington Drosophila Stock center | BDSC: 84589 |
| *5-HT2A-GAL4* | Bloomington Drosophila Stock center | BDSC: 84590 |
| *5-HT2B-GAL4* | Bloomington Drosophila Stock center | BDSC: 84591 |
| *5-HT7<sup>GAL4</sup>* | Bloomington Drosophila Stock center | BDSC: 86279 |
| *UAS-5-HT7<sup>RNAi</sup>* | Bloomington Drosophila Stock Center | BDSC: 27273 |
| *Dh44-GAL4* | Dr. Y. Kim | KDRC: 2543 |
| *Dh44R1<sup>Mi</sup>* | Dr. Y. Kim | BDSC: 23517 |
| *Dh44R2<sup>Mi</sup>* | Dr. Y. Kim | BDSC: 29129 |
| *Dh44-LexA* | Korea Drosophila Resource Center | KDRC: 2776 |
| *LexAop-mCherry* | Korea Drosophila Resource Center | KDRC: 1238 |
| *elav-GAL4* | Korea Drosophila Resource Center | KDRC 2690 |
| *Myo1A-GAL4* | Bloomington Drosophila Stock Center | BDSC 67057 |
| *UAS-mCD8::GFP* | Bloomington Drosophila Stock Center | BDSC 5137 |
| *UAS-mCD8::GFP* | Bloomington Drosophila Stock Center | BDSC 32184 |
| *UAS-Kir2.1* | Bloomington Drosophila Stock Center | BDSC 6596 |
| *tub-GAL80<sup>ts</sup>;UAS-Kir2.1* | Korea Drosophila Resource Center | KDRC 1103 |
| *UAS-dicer2* | Bloomington Drosophila Stock Center | BDSC 24645 |
| **Antibodies** |  |  |
| Mouse anti-GFP | Molecular probe | Cat # A11120 |
| Rabbit anti-DsRed | Takara | Cat # 632496 |
| Goat anti-mouse Alexa Fluor 488 | Thermo Fisher/Invitrogen | Cat # A11029 |
| Goat anti-rabbit Alexa Fluor 568 | Thermo fisher/Invitrogen | Cat # A11036 |
| **Chemicals, Enzymes and other reagents** |  |  |
| Agarose | Sigma-Aldrich Co. | Cat # A0224-050 |
| Sucrose | Sigma-Aldrich Co. | Cat # S0398-1KG |
| Brilliant blue | Walko Chemical Co. | Cat # 02712842 |
| Triton<sup>TM</sup> X-100 | Sigma-Aldrich Co. | Cat # X100-500ML |
| Phosphate buffer saline | Dyne Bio. | Cat # CBP3070 |
| Saponin | Junsei Chemical Co. | Cat # 31385-1510 |
| **Software** |  |  |
| *Origin Pro Version* | OriginLab corporation | RRID: SCR_002815<br>https://www.originlab.com |
| *Graphpad Prism* | GraphPad | RRID: SCR_002798<br>https://www.graphpad.com |

### Crop size scoring

Crop size scoring was performed as described previously.(Edgecomb et al., 1994) In brief, male flies were placed on food consisting of 1% agarose mixed with 5% sucrose and blue dye (brilliant blue FCF, 0.125 mg/mL; Wako Chemical Co., cat. # 027-12842) for 12 h. In total, 10–15 flies per group were dissected in 1X PBS under a microscope, and their dye-filled crops were then scored on a scale of 1–5 as follows:(Edgecomb et al., 1994) (1) the crop shrank and no dye was visible in the crop, (2) a small blue spot was visible in the crop, (3) the crop was wider and longer but extended toward the lateral side of the abdomen, (4) the crop was round and the abdomen was slightly swollen, and (5) the crop was maximally distended and the abdomen was extremely swollen. The percentages of flies with each crop score were calculated.

### Measurement of crop contraction

Crop contraction was measured as described previously, with some modifications.(Solari et al., 2017) In brief, male flies were fed 1% agarose mixed with 5% sucrose and blue dye overnight in a humidified chamber. The flies were cold anesthetized and fixed on a glass slide with glue. The fixed flies were then transferred and submerged in a Petri dish cover with 1X PBS (128 mM NaCl, 36 mM sucrose, 4 mM MgCl2, 2 mM KCl, 1.8 mM CaCl2, pH 7.1). The ventral cuticle was gently removed to view the crop. Subsequently, the number of crop contractions/min was manually counted using to stereomicroscope.

### Immunohistochemical analysis

For performing immunohistochemical analysis, freshly dissected brain samples were placed into an Eppendorf tube and intestinal samples were placed into a well of a 24-well tissue culture plate (Costar Corp.) on ice. The brain samples were fixed in 1 mL of 0.1% saponin with 1X PBS solution containing 4% paraformaldehyde, whereas the intestinal samples were fixed in 940 µL of 0.1% TritonX in 1X PBS solution containing 60 μL formaldehyde (37%) and incubated for 30 min. The brain samples were then washed thrice with wash buffer consisting of 1X PBS containing 0.1% saponin, while the intestinal samples were washed thrice with wash buffer consisting of 1X PBS containing 0.1% Triton X (15 min per wash). Subsequently, the samples were blocked with 1 mL of blocking buffer (1X PBS, 0.1% saponin, and 5 mg/mL BSA) at 4°C for 4 h. To perform immunostaining, the samples were incubated with the primary antibodies mouse anti-GFP (1:1,000, Molecular Probes, Cat # A11120) and rabbit anti-DsRed (1:1,000, Takara, Cat # 632496) at 4°C for 18 h. The samples were then washed thrice with wash buffer (15 min per wash) and treated with the secondary antibodies (1:200) goat anti-mouse Alexa Fluor 488 (Cat # A11029), and goat anti-rabbit Alexa 568 (Cat # A11036) at 4°C for 4 h. Finally, the samples were washed thrice and stored in 1.25X PDA buffer (187.5 mM NaCl, 37.5% glycerol, 62.5 mM Tris, pH 8.8) at 4°C for more than 1 h. The samples were then mounted and examined under a Leica Stellaris 5 confocal microscope.

### Statistics and reproducibility

*D*. *melanogaster* was selected as a model organism in this study. For the experiments, male flies were mostly used, unless specified otherwise. All the experiments were conducted under laboratory conditions. The appropriate number of replicates was established based on previous research. A sufficiently large sample size was used in all our assays to ensure that the results were representative and repeatable. No data points were excluded from the analysis. For each genotype, the data points indicate the values of individual replicates. The error bars in all the figures represent the standard errors of the mean (SEMs). The asterisks in the figures indicate statistical significance (\**P* < 0.05, \*\**P* < 0.01). All statistical analyses were performed using Origin Pro 8 software for Windows (ver. 8.0932, Origin Lab Corporation, USA).

## Author contributions

D.K.N. and S.D. conducted the experiments. D.K.N. analyzed the data, generated the figures, and wrote the original draft. Y.L. supervised the research, reviewed and edited the draft, and obtained the funding.

## Disclosure and competing interests statement

The authors have no competing interests.

## Acknowledgments

This work was supported by grants to Y.L. from the National Research Foundation of Korea (NRF) funded by the Korean government (MIST) (RS-2021-NR058319) and the Biomaterials Specialized Graduate Program through the Korea Environmental Industry & Technology Institute (KEITI) funded by the Ministry of Environment (MOE). D.K.N. and S.D. were supported by the Global Scholarship Program for Foreign Graduate Students at Kookmin University in Korea.

## Supplementary material

**EXPANDED VIEW FIGURE PDF**

